# Numerical Simulation of population development using the function of the age distribution of birthrate

**DOI:** 10.1101/031047

**Authors:** Alexandr N. Tetearing

**Affiliations:** Saint Petersburg State University, Russia

## Abstract

Numerical model of population behaviour under conditions of full-fed existence of mortal persons in the personal areas, simulated with use the function of age distribution of birth rate in the form of normal (gaussian) distribution. Graphs of historical changes in the function of age distribution of population.

## Simulation algorithm

The knowledge of the function of age distribution of birth rate *Bg*(*t*) allows us more accurately simulate the process of population development. Especially under conditions of the limited food resource. Because, in this simulation, we can more exactly calculate the form of function of the age distribution of population mass *Ag*(*t*) at each time point (more precisely for each step of the calculations).

Herewith, the each age group receives and spends an energy, and produces a posterity with different relative intensity, and these age groups differ from each other according to these indicators.

Besides, in the population development process, at certain moments of time, the calculated mass of certain age groups can be a negative value – for example, when the energy, that the group receives from external ambience, is less than the current energy expenses of this group. The algorithms of numerical simulation do not always allow us to take into account the emergence of age groups with negative mass in the population. The below-described algorithm of step-by-step numerical simulation of population development allows us to easily keep track of such moments of time and correct the current age distribution of mass by replacing of a negative mass of age group by zero value. That is to say, herewith we believe that this age group is dead and the mass of this group is not negative, but it is equal to zero.

The algorithm of numerical simulation of the process of change in population mass can be as follows.

We choose the step of calculations Δ*t.* Herewith, the function of age distribution of population mass contains of *n* age groups, where 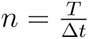. Here *T* is the life time of persons in population.

We denote the mass of the age group with number *n* as *m_n_*.

At each time point, we have to calculate (using the known laws of the development of population under different conditions of environment) the amount of energy *E_n_,* that the group receives from environment during the time Δ*t,* the energy costs 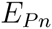, that the group spends on the maintenance of the population mass (during the time Δ*t*, the energy costs 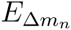 for the birth of a posterity (during the time Δ*t*, and the increase of mass 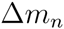 of newborn posterity, which this group produces during the time Δ*t.*

The integer index *n* means here the serial number of age group.

We assume, that at initial moment of time *t* = 0 the population consists of one person with zero age and mass *m*_0_. We denote the total mass of population at moment of time *t* = 0 as *M*_1_ or *M*(0). That is to say:

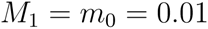

We denote as *N_n_* the amount of persons in the group with index *n*.

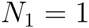

The mass of newborn posterity 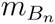, that the age group with index *n* produces, is calculated by the formula:

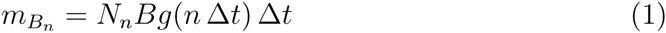

Here *Bg*(*n*Δ*t*) is the value of function *Bg*(*t*) at moment of time *n*Δ*t.* Naturally, the smaller the step of calculations Δ*t,* the more accurate this formula is.

To find the total mass *M_B_* of newborn posterity, that the population produces for a time Δ*t,* we need to summarize the masses of posterity, that every age group produces:

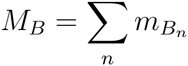

In our calculations, we assume that the number of newborn cells (the first cells of future organisms) does not depend on the conditions of existence of female. But, if the female lives under conditions of hungry existence, then her cub (which develops a certain time in the womb of the female and consists already of ensemble of the biological cells) has a smaller mass at birth, in contrast with mass of cub of female, which lives under conditions of profusion of food.

Of course, the biologists can use in their calculations the own assumptions about conditions of existence of population (which differ from the aforementioned conditions). We show here only a certain base algorithm of the calculation of function of the population development, and it is one of the set of possible algorithms.

We build the graph of the development of population as a plurality of discrete points with step Δ*t*. The smaller Δ*t* is, the more exactly a solution. But, generally speaking, the step-by-step program algorithms, like this, accumulate progressive error of calculations (the final inaccuracy is proportional of the number of steps), and the computation time is inversely proportional to the value Δ*t*. So, we can define the reasonable criterion for choice of the computation step as follows. If, with reduction of computation step (for example, with reduction by half), the form of the graph of solution will change little (the form is visually unchanged), we can conclude that the step is small enough.

In each step of the calculation, we have to find the amount of energy *E_n_*, that each age group receives from the environment:

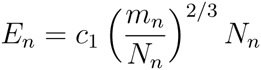

The proportionality factor *c*_1_ is here a certain constant, *m_n_* is the mass of the age group, *N_n_* is number of persons in the group, 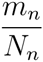 is the mass of one person in this group.

Here we believe that the amount of energy *E_n_*, which the age group with index *n* receives from the environment, is proportional to the surface area of person’s body.^1^

Since, ultimately, we aim to obtain the function, describing the change in the population mass, we need to transform all the energy revenues to the population mass. We assume that, in absence of other expenses, the energy, which the organism receives from environment, can be converted by organism to the additional mass of the body (for example, the energy is transformed by organism to the muscular mass or adipose tissue):

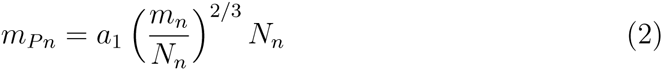

Here 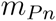 is an additional mass, that can be received from environment for time Δ*t* by age group, which has the index *n*, the mass *m_n_* and the number of persons *N_n_*. The factor *a*_1_ is a certain constant.

Thus, we consider that the population lives the full-fed existence and consumes as much food resource as can be eaten (roughly speaking, the amount of food, that one person consumes, is proportional to the area of person’s stomach).

At each step of the calculation, we have to take into account also the energy expenses of age group, that persons spend on the maintenance of their organisms mass. It is logical to believe that these expenses 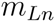 (the expenses of energy or adipose tissue) are proportional to the mass of age group *m_n_* and length of time interval Δ*t:*

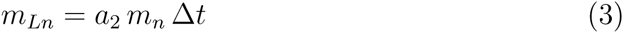

We also believe that the person’s expenses on the mining of energy from environment are also proportional to the mass of organism. Thus, in this case, we believe that these expenses are already taken into account in the coefficient *a*_2_.

We need also to take into account (as expenses of the mass) the expenses of the mass 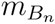, that the age group spends on birth of posterity. Thus, we can write, for each age group with index *n*, the following equation:

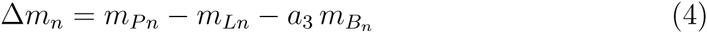

Here 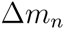 is increment of the mass of the age group with index *n* during time interval Δ*t;*

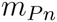 is the mass of organism (for example, the adipose tissue or muscular mass), that can be received by organism from energy, which the age group receives from environment for time Δ*t*;

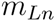 are expenses of the age group on energy maintenance of own mass;

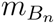 is the mass of newborn posterity, which the age group with index *n* produces for time interval Δ*t*;

*a*_3_ is a certain constant, which is always greater than unity, because, at production of posterity, the mass of the parental organism can not be converted to the mass of daughter organism without a some energy losses.

The equation (4) is the equation of life (the equation of the energy balance), which is written for separate age group. We write the equation using no energy units, but a units of mass.

In numerical solving this equations – when the constants, that we use in the algorithm, can be set quite arbitrarily – we need, on each step of calculations, to check the following condition:

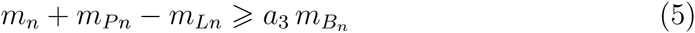

This condition reflects the fact that the organism can not for time Δ*t* produce the posterity more, than the sum of organism mass and mass, that organism received from environment (with taking into account the expenses on servicing of own mass). If this condition is violated, we need to adjust the algorithm.

At correction of algorithm, we must take into account that the mass of age group can not be negative. If, when computing, the mass of group *m_n_* is negative, we will consider that the corresponding group ceased its existence (at following step of calculations, the current mass of this group will be equal to zero).

If the mass of the age group is not enough to produce the posterity in full (with amount, that the function of the distribution of birth rate dictates), this group produces fewer cubs, that corresponds to the actual mass of the group.

Moreover, the computing algorithm may also include some restrictions on the production of posterity.

Usually, for highly developed organisms the whole mass of parental person can not be converted into mass of cells of the future posterity, but, in our simplest algorithm, this is quite possible.

The final algorithm will describe the development of population, where the persons live in the personal areas under conditions of full-fed existence.

## Results and graphs

The graph of the numerical solution, that was obtained as a result of running of this simulation algorithm, is shown in figure 1.

**Figure 1:**
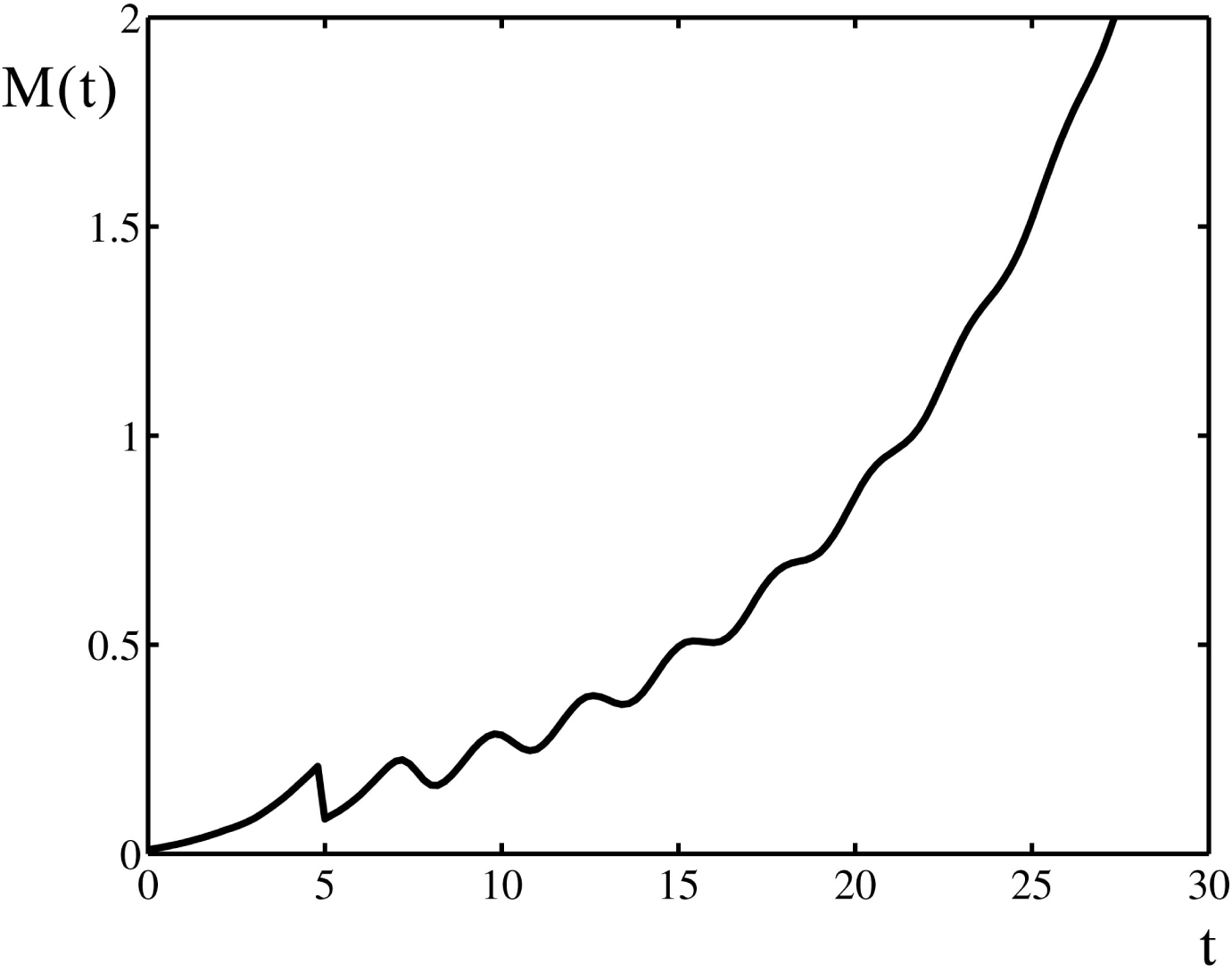
Graph of function of change in population mass obtained as result of the numerical simulation with use of known function of the age distribution of birth rate *Bg*(*t*).

In the calculations, we used the following constants: the time of life of persons is *T* = 5; the calculation step is Δ*t* = 0.2; the constants are *a_1_* = 0.34, *a*_2_ = 0.32, *a*_3_ = 1.5; the initial mass of population (the mass of the first person) is *m*_0_ = 0.01.

In the initial time interval, from *t* = 0 to *t* = 5, the graph of function in figure 1 is increasing exponential function. At moment of time *t* = 5 the first deaths of persons from old age occur in population. After this point the shape of function *M*(*t*) is changed.

For plotting of the graph in figure 1, as the function of the age distribution of birth rate, we used the following (a priori known) function:

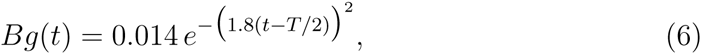

that is shown in figure 2.

**Figure 2:**
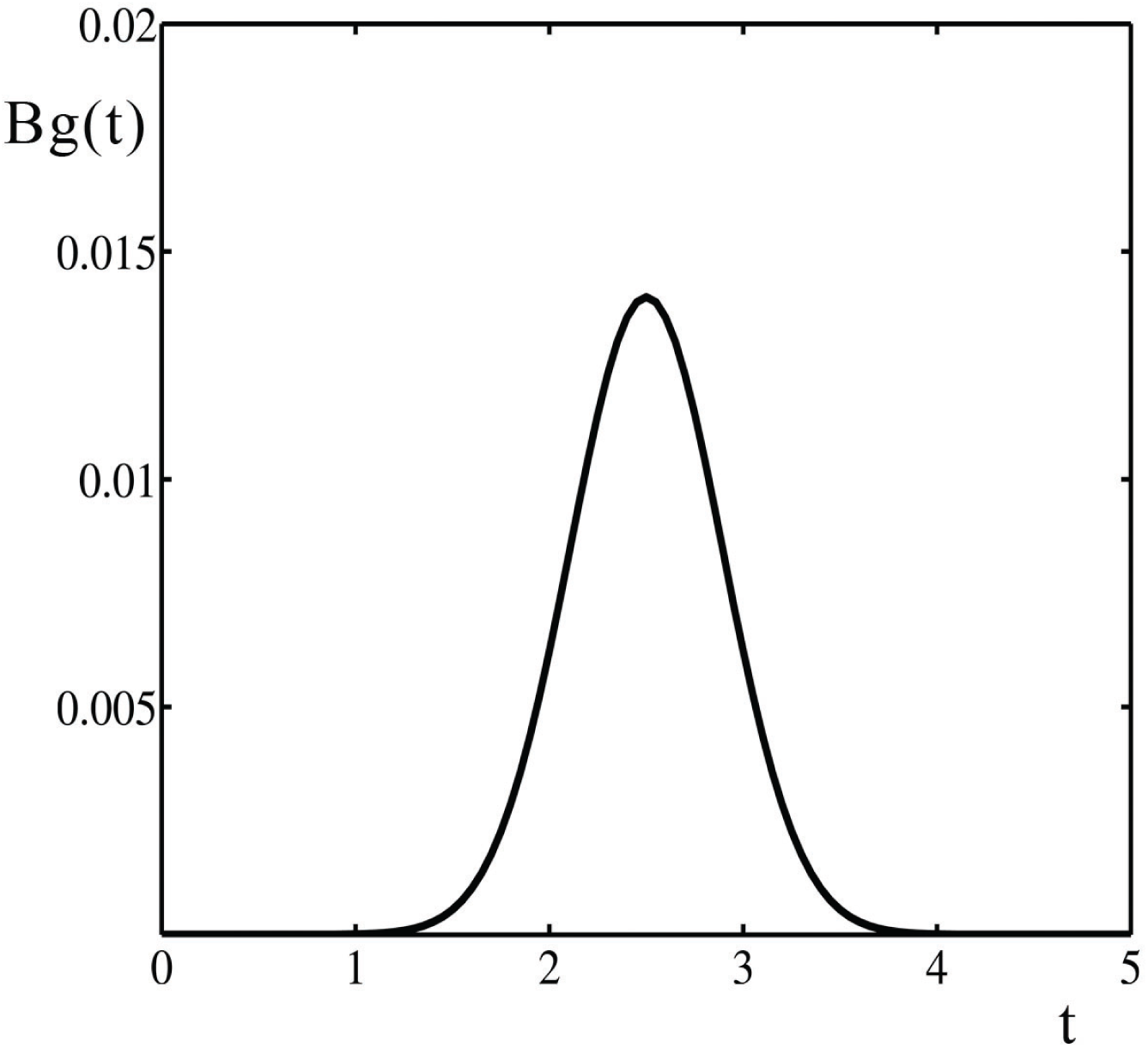
Graph of known function of the age distribution of birth rate.

The numerical calculations give us a good opportunity to see not only the changes in population mass, but also the changes, which occur, with development of population, in its age composition.

In figure 3 we see the graph of the time history of function *Agd*(*t*), where *Agd*(*t*) is the function of the age distribution of population mass.

**Figure 3:**
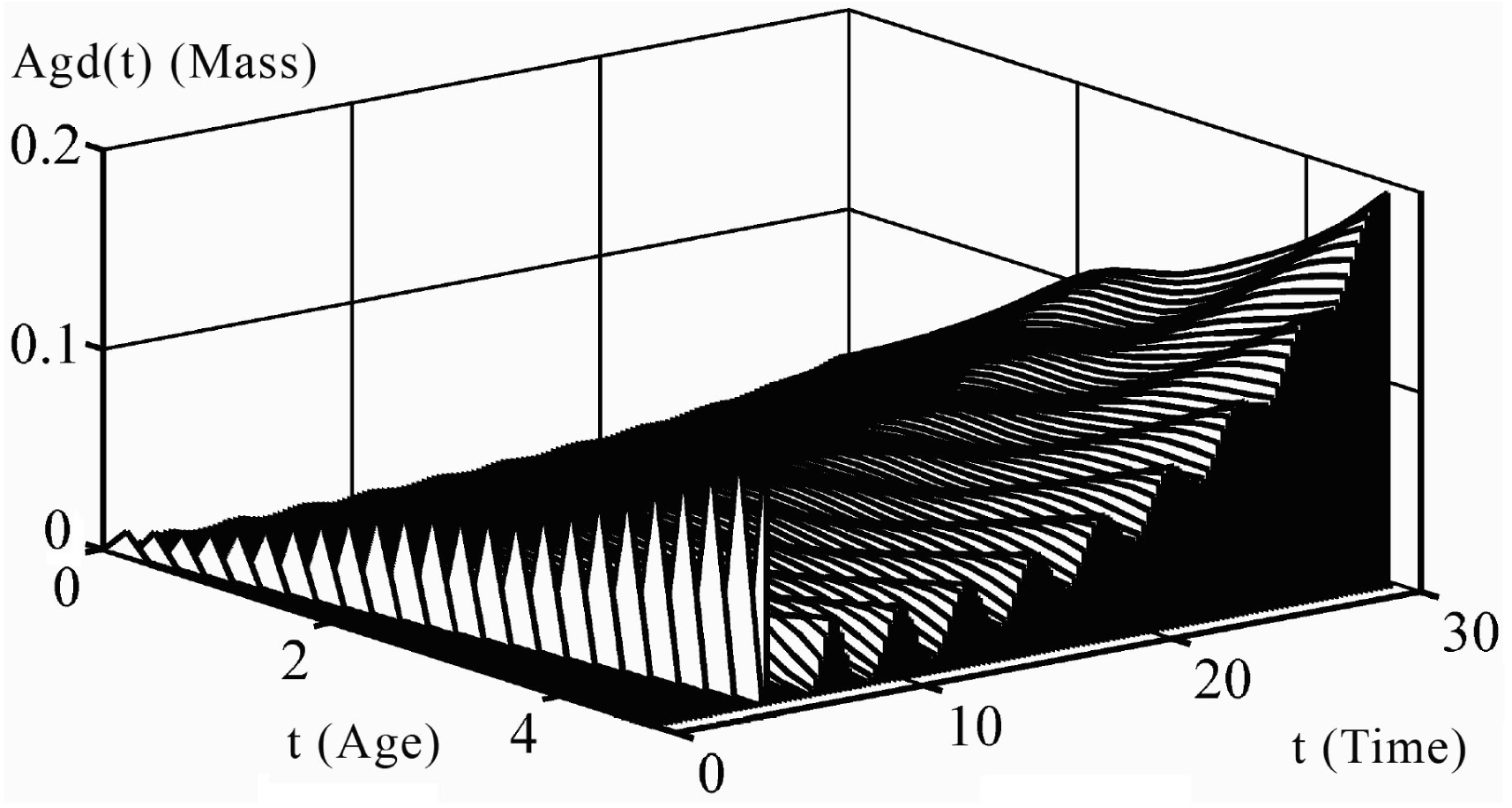
Graph of the change in the function of age distribution of the population mass.

The horizontal axis “Time”, that is directed to the right (from *t* = 0 to *t* = 30), is a current time. The horizontal axis “Age”, that is directed to the left (from *t* = 5 to *t* = 0), is the age of persons. The vertical axis “Mass”is the mass of the group of persons with age, that is specified on the “Age” axis, at the corresponding moment of time, that is specified on the “Time” axis.

For clarity, the separate figure 4 shows the temporal cross sections of this three-dimensional surface, that were made with a vertical cutting planes, which are parallel with the axises *Age* and *Agd,* at moments of time *t* = 10, *t* = 20, and *t* = 30.

**Figure 4:**
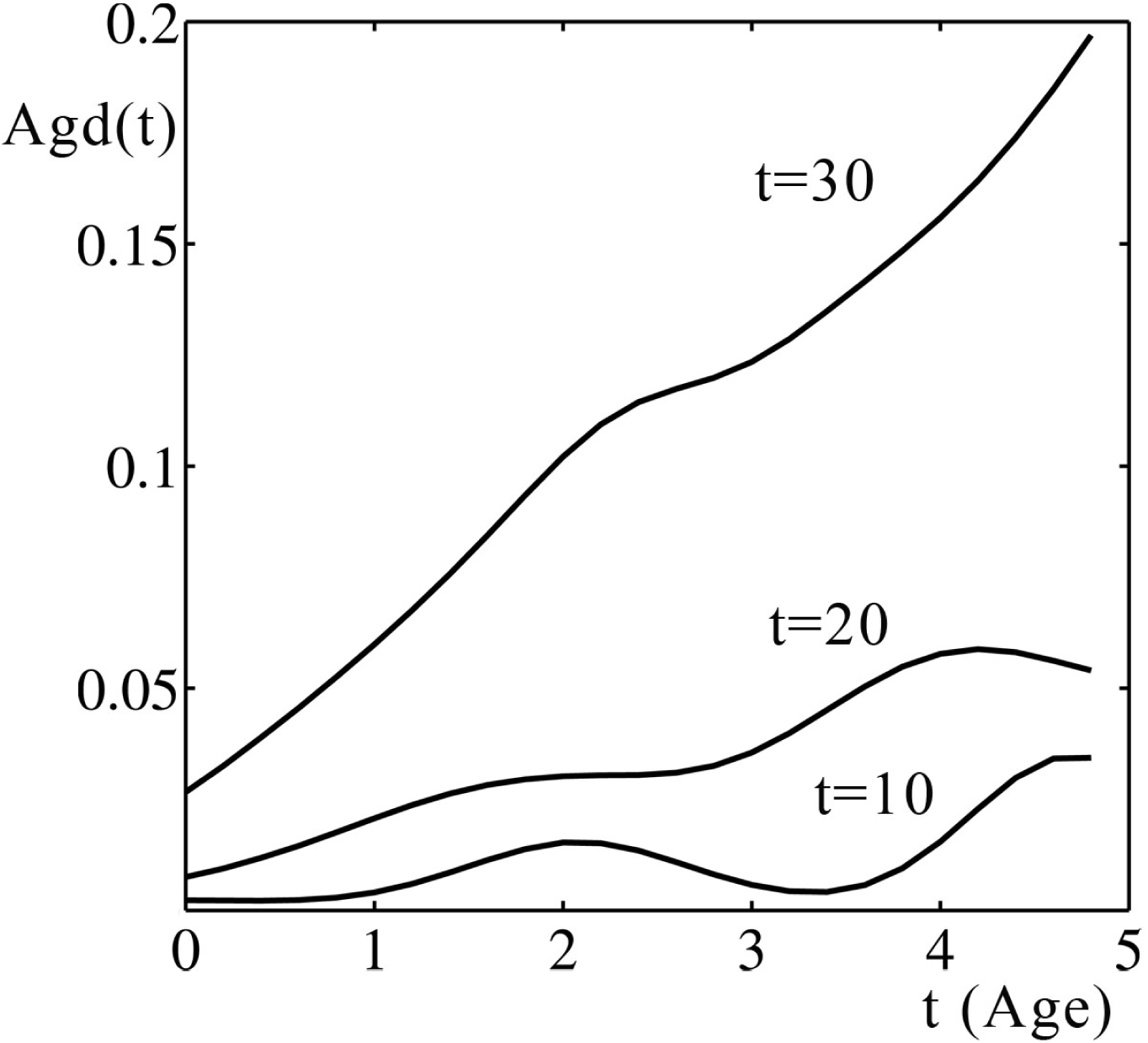
Functions of the age distribution of mass of population, which are obtained at different time points.

Since the distribution of persons in age groups is discrete (the population is separated into groups, where the age difference in each group is Δ*t* = 0.2), the each graph a little bit not reached the right edge of the picture (there is the gap of 0.2 units). That is to say, there are persons aged 4.8 in the population (if the time is measured in years), but there are no persons aged 5 – we consider that these persons have already died from old age.

The figures 3 and 4 show that the great part of the population mass consists of elderly persons – the mass of young persons is small, but the mass of old persons is large.

But if we look at an numerical distribution of persons in population instead of the mass distribution (this numerical distribution is shown in figure 5), we see that, with large values *t*, the population develops so that the young persons dominate in population – the number of young persons in population larger, than the number of elderly persons.

**Figure 5:**
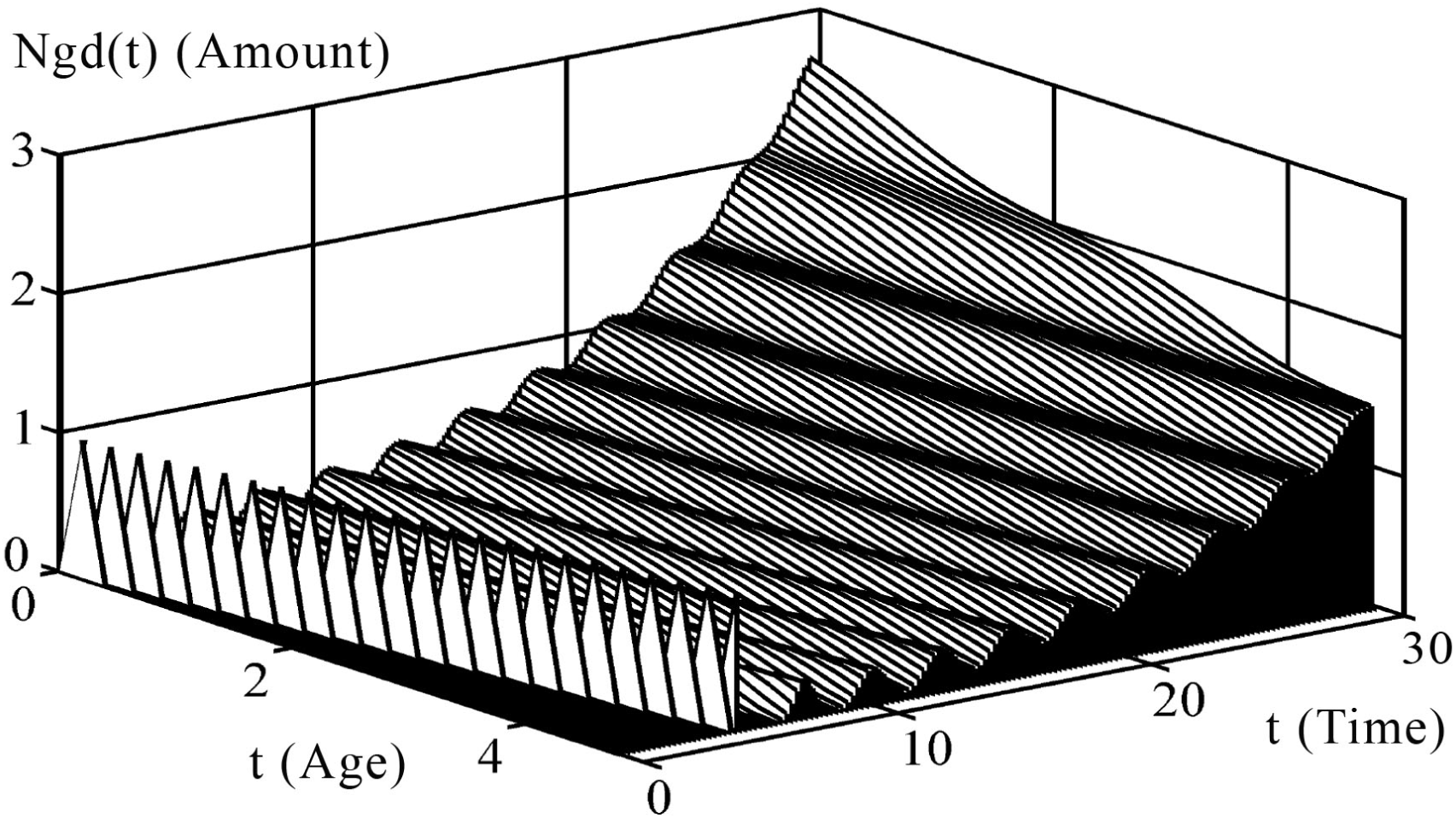
Graph of the change in the function of age distribution of the number of persons.

The vertical temporal cross sections of the three-dimensional surface from the figure 5 are shown in figure 6, where the functions of the age distribution of number of persons are shown at moments of time *t* = 10, *t* = 20 and *t* = 30.

**Figure 6:**
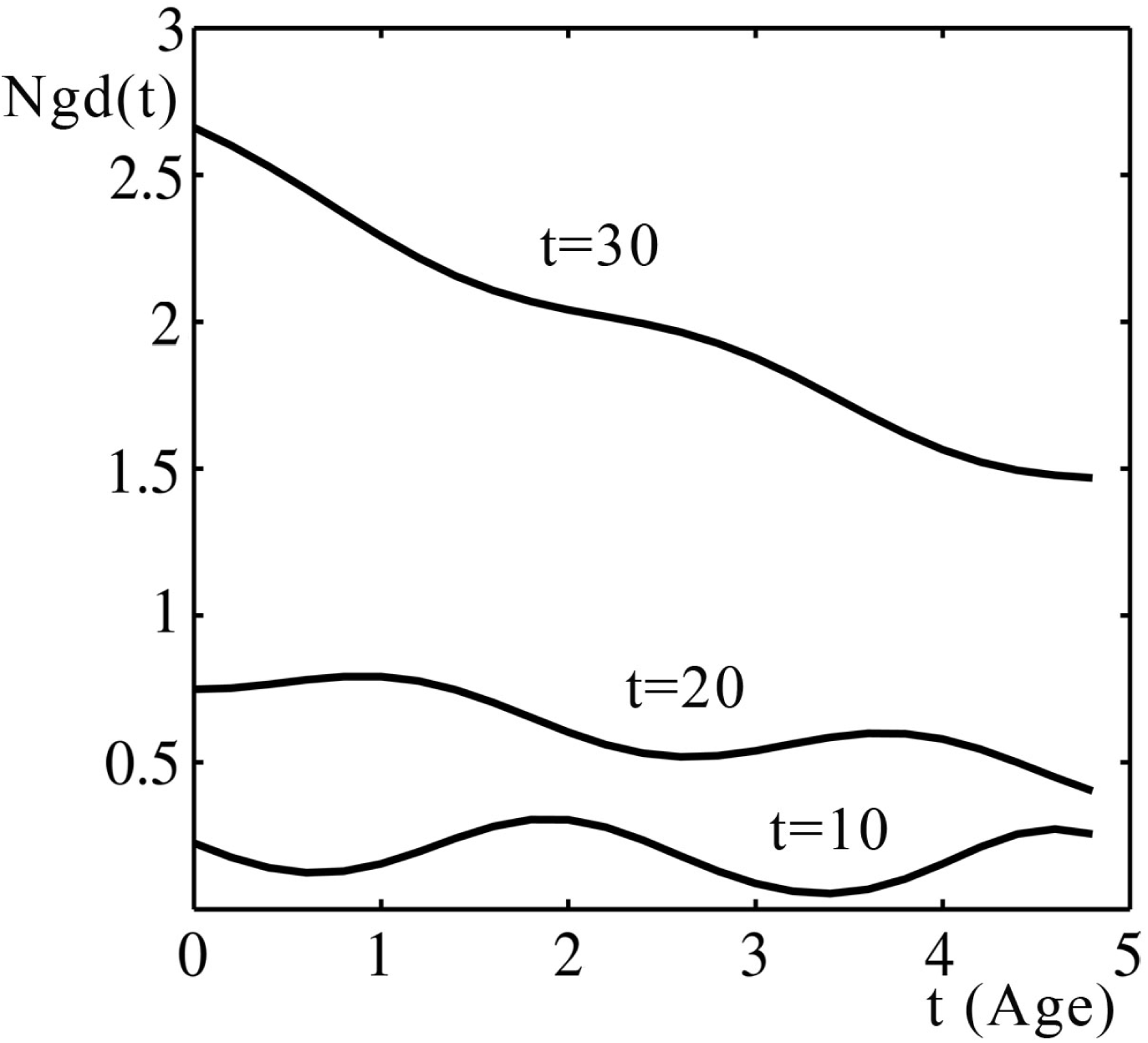
Functions of the age distribution of number of persons in population, which are obtained at different time points.

## Discussion of results

The function, shown in figure 1, can be described by the following equation:

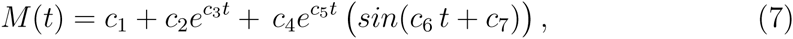

where *M*(*t*) is a total mass of population, *c_i_* are some constants.

This function is the solution of the equation of life for the population which develops under conditions of full-fed existence of persons in the personal areas with taking into account the mortality of persons in population.

The development of population under these conditions is described by the following equation:

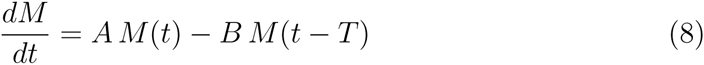

Here *A* and *B* are the certain constants, *T* is the life time of persons in population. The equation has a solution in the form of exponential function:

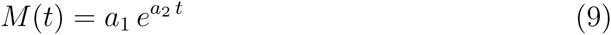

We can write the formula for function of numerical age distribution *Ng*(*t*), corresponding to these equations:

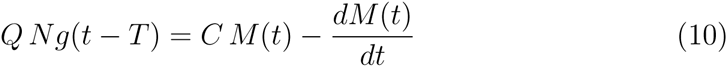

Since the derivative of exponential function is an exponent, we have:

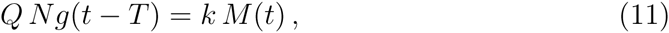

where *k* is a certain constant factor.

It means that, in practical calculations, we can use the equations (8) as exact equations (not as approximate equations), when we study the decision *M*(*t*), which is the exponential function (or the sum of exponents). We assume here that the function of numerical age distribution is an exponent in the form:

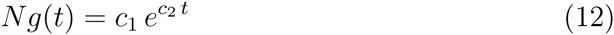

Then, by definition:

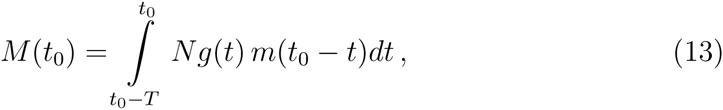

or:

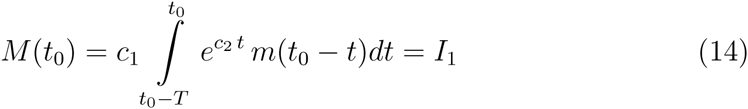

Here *m*(*t*) is a function of growth of organism mass, *I*_1_ is the our designation of this integral. If we write the function *M*(*t*) in the point *t*_0_ + *x*, where *x* is a certain fixed increment of the argument.

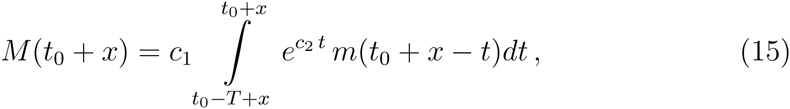

the formula is transformed as follows:

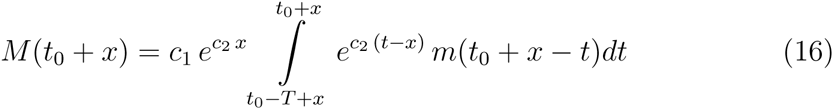

Because, in the integration, the variable *t* goes through the same values, as in the formula (14), consequently, we have:

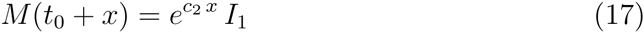

Here we use the following property of a definite integral of function with a delayed variable:

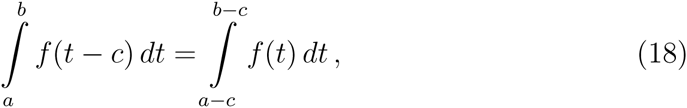

where *c* is a free constant.

If we rewrite in the same manner the formula for *M*(*t*_0_ + 2*x*), we find that:

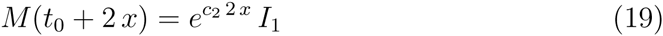

or, in general:

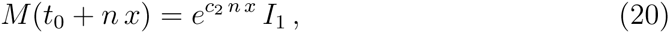

where *n* is a positive integer.

If we choose the timeline so that *t*_0_ is equal to zero, we obtain the formula:

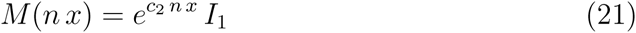

Consequently, we can write:

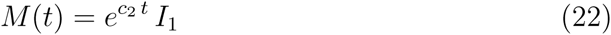

where *I*_1_ is a certain constant:

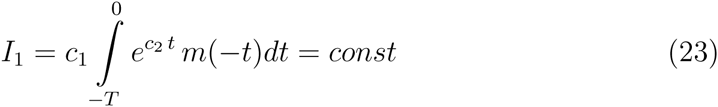

Thus, if the function of the change in population mass *M*(*t*) is an exponent in the form 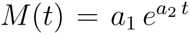, then its numerical age distribution *Ng*(*t*) is also an exponential function (regardless of the form of the function of growth of organism mass *m*(*t*), provided that the form of function of growth of organism mass does not change with time).

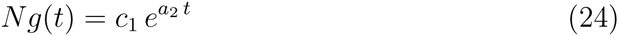

The constant *c*_1_ is defined by formula:

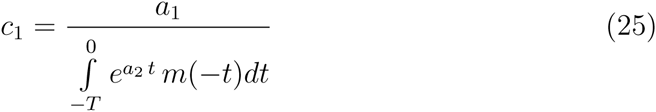

If the function *M*(*t*) is a sum of exponents:

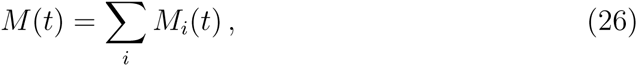

then (by the additivity property of integration procedure) we can calculate the function of the numerical age distribution *Ng_i_*(*t*) for each exponent and summarize the results to get the general function:

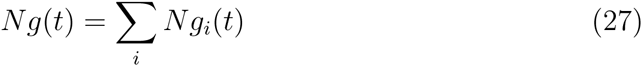

Looking at figure 1 we observe that, in course of time, when the sinusoidal oscillations of the function *M*(*t*) are damped out, the age distribution of number of persons in population *Ngd*(*t*) becomes more and more like a decreasing exponential function. We showed it above that, for population with exponential development, the age distribution of number of persons in population (when the function of the growth does not change with time) is also an exponential function.

Thereby, we can conclude that the aforedescribed algorithm of the numerical simulation of process of population development demonstrates the results, which are quite comparable with solutions of the simplest equations of life of biological populations.

## Footnotes

1 As a the first approximation, the area of body surface is proportional to the body volume to the power 2/3. The volume of the body is proportional to the mass.

